# Energetics based epitope screening in SARS CoV-2 (COVID 19) spike glycoprotein by Immuno-informatic analysis aiming to a suitable vaccine development

**DOI:** 10.1101/2020.04.02.021725

**Authors:** Amrita Banerjee, Dipannita Santra, Smarajit Maiti

## Abstract

The recent outbreak by SARS-CoV-2 has generated a chaos in global health and economy and claimed/infected a large number of lives. Closely resembling with SARS CoV, the present strain has manifested exceptionally higher degree of spreadability, virulence and stability possibly due to some unidentified mutations. The viral spike glycoprotein is very likely to interact with host Angiotensin-Converting Enzyme 2 (ACE2) and transmits its genetic materials and hijacks host machinery with extreme fidelity for self propagation. Few attempts have been made to develop a suitable vaccine or ACE2 blocker or virus-receptor inhibitor within this short period of time. Here, attempt was taken to develop some therapeutic and vaccination strategies with a comparison of spike glycoproteins among SARS-CoV, MERS-CoV and the SARS-CoV-2. We verified their structure quality (SWISS-MODEL, Phyre2, Pymol) topology (ProFunc), motifs (MEME Suite, GLAM2Scan), gene ontology based conserved domain (InterPro database) and screened several epitopes (SVMTrip) of SARS CoV-2 based on their energetics, IC50 and antigenicity with regard to their possible glycosylation and MHC/paratopic binding (Vaxigen v2.0, HawkDock, ZDOCK Server) effects. We screened here few pairs of spike protein epitopic regions and selected their energetic, IC50, MHC II reactivity and found some of those to be very good target for vaccination. A possible role of glycosylation on epitopic region showed profound effects on epitopic recognition. The present work might be helpful for the urgent development of a suitable vaccination regimen against SARS CoV-2.

## Introduction

An outbreak of a novel Coronavirus, Severe Acute Respiratory Syndrome CoV-2 (or SARS CoV-2 or COVID-19) infection is threatening the humanity, globally occurring from last week of December 2019. As a result, a massive loss of human health status and global economy are becoming unaccountable. As of current situation, SARS CoV-2 claimed more than 40,000 lives from more than 800,000 infected persons globally [1]. The outbreak started from the Wuhan province of China and spread more than 195 countries with most adverse effects in China, Italy, Iran, Spain, the United States, France, Germany, Britain and several other countries. Any type of therapeutic strategies starting from the blocking of viral entry, inhibition of spike proteins association with host ACE-2 (angiotensin converting enzyme type 2), modulations of interfering kinase activity, inactivation of viral genome expression-packaging and vaccination against this virus is the demand of the present situation. Regarding the vaccination strategies, it is assumed that frequent mutation results in anomalies in its surface/ spike proteins [2,3]. Mostly resembling the features of SARS CoV global outbreak (2003, https://www.who.int/csr/sars/en/), this virus unlikely manifested it’s extremely high grade of virulence, spreading capability and stability across the geographical barrier (or specifically colder place, aged persons or specific genders; yet to be clarified) [4].

The positive selective pressure could account for the stability and some clinical features of this virus compared with SARS and Bat SARS-like CoV [5]. Stabilizing mutation falling in the endosome-associated-protein-like domain of the nsp2 protein could account for COVID-2019 high ability of contagious, while the destabilizing mutation in nsp3 proteins could suggest a potential mechanism differentiating COVID-2019 from SARS CoV [5]. Nevertheless, nutritional and immunological statuses are also important factors for the screening of the therapeutic strategies for the affected and sensitive persons. Possible medications or immunizations from the existing drugs or infusion of convalescent plasma should be conducted with utmost care to the COVID 19 patients [6]. Advanced precautionary steps and therapeutic interventions should be formulated taking into account of several personal and community factors [7]. Development of a successful and reproducible vaccination protocol and its human trial may take longer time for the issues of mutation and large number glycan shield and epitope masking on the SARS CoV 2 proteins [8].

In a series of medication regimen, 1 (AT1R) blockers is used for reducing the severity and mortality from SARS-CoV-2 virus infections [9]. Chloroquine and Hydroxychloroquine are now being prescribed somewhere to fight COVID-19 for the time being [10,11]. Human coronaviruses and other influenza viruses resulted in epidemic in last 2 decade in different parts of the world. The anomalies between severity and spreading between the origin site, China and the other parts of the World (European and North America countries) might have some indication. Common human CoVs may have annual peaks of circulation in winter months in the US, and individual human CoVs may show variable circulation from year to year. [12]

Colder climate and prior exposure to other human coronaviruses, or influenza or flu viruses or possible vaccination against those might develop antibody dependent enhancement (ADE) of immunological responses during recent SARS CoV-2 exposure. ADE might have modulated immune response and could elicit sustained inflammation, lymphopenia, and/or cytokine storm [13,14]. Possibly, that could be one of the reasons (more history of exposure with CoVs beside weaker immune system) for older people being more affected by the present SARS CoV-2. Moreover, both helper T cells and suppressor T cells in patients with COVID-19 were below normal levels. The novel coronavirus might mainly act on lymphocytes, especially T lymphocytes [15]. Strong inflammatory events could be the initiator of the collapsing environment during COVID-19 infection. In most of the death cases in COVID-19 infections, acute respiratory failure is followed by other organs like kidney anomalies. In these cases inflammatory outburst might have worsened the infection and post viral-incubation situations [16,17]. Recent studies in experimentally infected animal strongly suggest a crucial role for virus-induced immunopathological events in causing fatal pneumonia after human CoV infections [18]. So, combined anti-viral and anti-inflammatory treatment might be beneficial in these cases [19]. SARS-based available immune-therapeutic and prophylactic modalities revealed poor efficacy to neutralize and protect from infection by targeting the novel spike protein. [20].

In this background, critical screening of the spike sequence and structure from SARS CoV-2 by energetic and IC50 based immunoinfrmatics analysis may help to develop a suitable vaccine. So, in the current study we were intended to analyze the spike proteins of SARS CoV, MERS CoV and SARS CoV 2 and four other earlier out-breaking human corona virus strains. We critically compared SARS CoV and SARS CoV 2 spike-proteins, domains, motifs and screened several epitopes based on their energetics, IC50 and antigenicity employing several bio/immuuno –informatics software with regard to their possible glycosylation and MHC/paratopic binding effects. The present work might be helpful for the urgent development of a suitable vaccination regimen.

## Materials and Methods

### Sequence retrieval

The spike glycoprotein sequences of four human coronavirus (HKU1, NL63, 229E and OC43), MARS Coronavirus (NC_038294.1:21455-25516), SERS Coronavirus (NC_004718.3:21492-25259) were retrieved from viruSITE: integrated database for viral genomics [21], and SARS coronavirus 2 isolate Wuhan-Hu-1 (COVID 19) was retrieved from *National Center for Biotechnology Information* (NCBI) biological database (https://www.ncbi.nlm.nih.gov/).

### Structure Prediction and Structure Quality Assessment

Tertiary structures of selected coronavirus (CoV) spike proteins were predicted/ validated using Phyre2, Protein Homology/analogy Recognition Engine V 2.0 [22] and SWISS-MODEL [23]. In Phyre2 structures were predicted against 100,000 experimentally designed protein folds. Predicted structures were subjected to analysis in SWISS-MODEL for QMEAN Z-score calculation which includes cumulative Z-score of Cβ, All atoms, Solvation and Torsion values prediction. RAMPAGE: Ramachandran Plot Analysis server [24] was used for protein 3D structures quality assessment. The summation of number of residues in favored regions and in additionally allowed regions was considered for percent (%) quality assessment.

### Protein Structural Alignment

Predicted tertiary structures were visualized and aligned using PyMol molecular visualization system. Pymol assigns the secondary structure using a secondary structure alignment algorithm called “dss”, where the sequences of two structures were aligned first then the structures were aligned. For the visualization of molecules a high-speed ray-tracer molecular graphics system was used.

### Secondary structure analysis

Secondary structural analysis and their 3D folding patterns were analyzed in the form of topology using ProFunc; a protein function predicting server using protein 3D structures [25]. In protein classification, topology analysis plays an independent and effective alternative to traditional structural prediction. Topological differences between two structures indicated differences in protein folding and flexibility.

### Sequence Comparison

Sequence comparisons among selected CoV spike glycoproteins were conducted through multiple sequence alignment using Clustal X2 [26]. Conserved motifs were identified using MEME Suite (http://meme.sdsc.edu/meme/cgi-bin/mast.cgi) server. MEME Suite represents the ungapped conserved sequences which are frequently present in a group of related sequences. The 7 motif number has been defined in the current study for motif finding. Whereas, GLAM2Scan tools was used for the identification of gapped motifs within the related sequences. Conserved motifs were represented through LOGO using GLAM2Scan tools of MEME Suite server. Identified motifs were subjected to annotation using protein BLAST (https://blast.ncbi.nlm.nih.gov/Blast.cgi?PROGRAM=blastp&PAGE_TYPE=BlastSearch&LINK_LOC=blasthome) and finally functional gene ontology based conserved domain identification was conducted using InterPro: Classification of protein families interactive database [27].

### Epitope Designing

Conserved epitopes of SARS Cov-2 spike glycoprotein were identified using SVMTrip: A tool which predicts Linear Antigenic Epitopes [28]. SVMTrip predicts the linier antigenic epitopes by feeding Support Vector Machine with the Tri-peptide similarity and Propensity scores of different pre-analyzed epitope data. Annotation of predicted epitopes was performed through protein BLAST. SVMTrip have gained 80.1% sensitivity and 55.2% precision value with five fold cross-validation. For epitope prediction 20 amino acid lengths was selected.

### Analysis for Epitopes binding efficiency to MHC Class II

The Major Histocompatibility Complex (MHC) binding efficiency of predicted epitops was performed using Immune Epitope Database (IEDB) and Analysis Resource [29]. A total of 5 DPA, 6 DQA and 662 DRB alleles from MHC class II were screened for the detection of best interactive alleles on the basis of highest consensus percentile rank and lowest IC50 value. All the analyses were performed on Human Class II allele, using frequently occurring alleles (frequency > 1%), peptide length of 9mers was selected; consensus percentile rank ≤ 1 was used for the selection of peptides.

### Antigenecity Prediction

Antigenecity of predicted epitopes were determined using Vaxigen v2.0 protective antigen, tumour antigens and subunit vaccines prediction server [30]. Vaxigen v2.0 uses auto cross covariance (ACC) transformation of selected protein sequences based on unique amino acid properties. Each sequence was used to find out 100 known antigen and 100 non-antigens. The identified sequences were tested for antigenecity by leave-one-out cross-validation and overall external validation. The prediction accuracy was up to 89%.

### Molecular Docking

The structure of MHC class II HLA-DRA, DRB molecule (PDB ID: 2q6w, 5jlz) and fully glycosylated COVID 19 spike protein structure (PDB ID: 6svb) was retrieved from Protein Data Bank (PDB) and Docking was performed using HawkDock [31] and ZDOCK [32] Server generating 100 docking solutions. Among them best 10 were analyzed based on docking scores and binding free energy value calculation.

## Results and Discussion

### Structure Prediction and Structure Quality Assessment

The spike glycoprotein structures of seven coronavirus including the recent outbreaking strain SARS CoV-2 (Covid 19) presented in table 1. The Ramachandran plot data and structural alignment data suggests that SARS CoV and SARS CoV-2 (Covid-2) has higher degree of alignment (Table 1). The protein sequences were ranged from 1173-1356 amino acids. An effort was made for epitope based peptide vaccine development by searching MHC-I and II classes compatible sites and the results yet to come [33]. In the current study, energetic and Inhibition Concentration50 based selection of SARS CoV-2 spike epitope and its possible glycosylation effect/structural-hindrance have been evaluated. This may help in urgent vaccination strategies in the current disastrous situation.

**Table 1.**
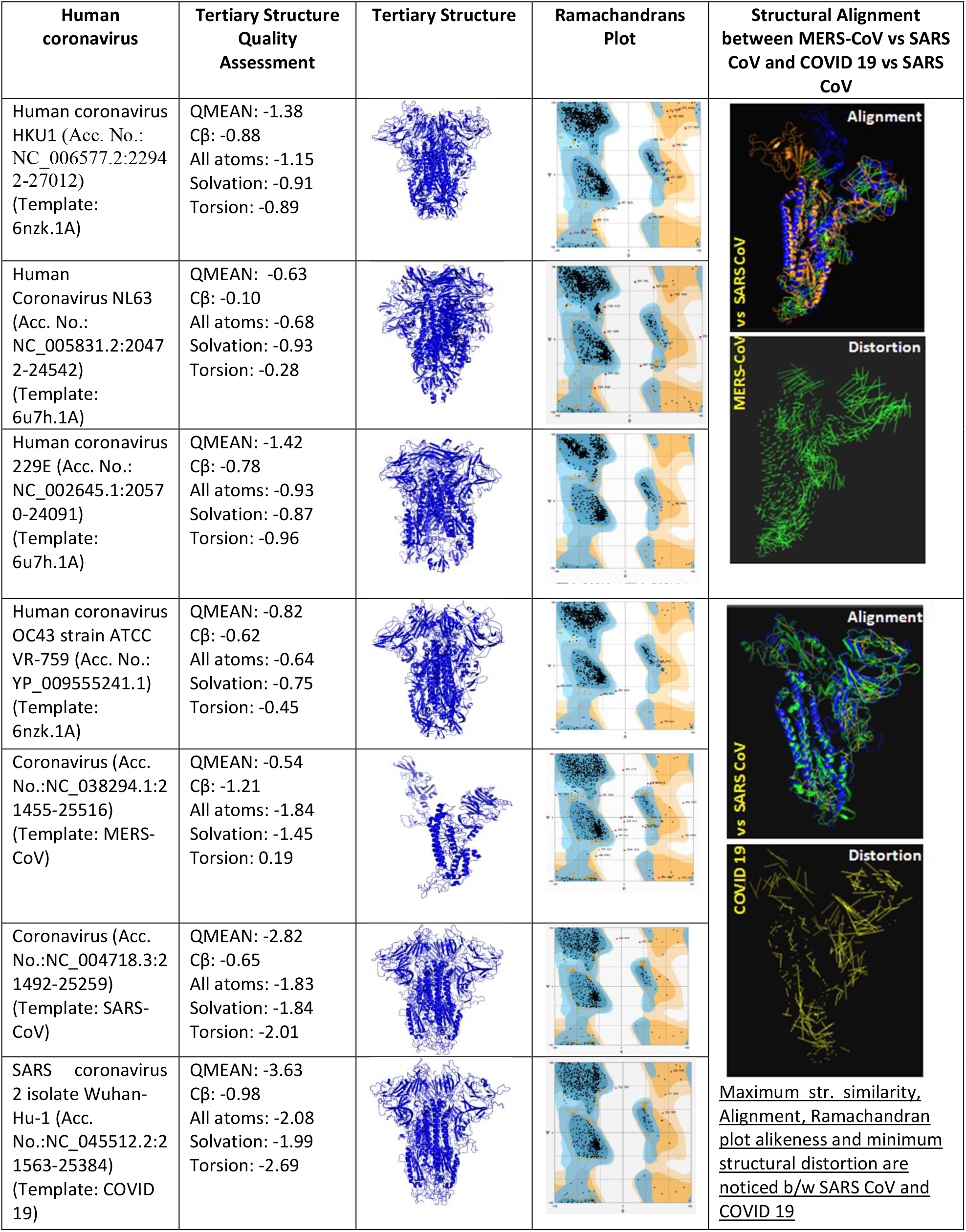
Global quality estimates of different coronavirus spike glycoproteins.

| Human coronavirus | Tertiary Structure Quality Assessment | Tertiary Structure | Ramachandrans Plot | Structural Alignment between MERS-CoV vs SARS CoV and COVID 19 vs SARS CoV |
| --- | --- | --- | --- | --- |
| Human coronavirus HKU1 (Acc. No.: NC_006577.2:22942-27012) (Template: 6nzk.1A) | QMEAN: -1.38<br>Cβ: -0.88<br>All atoms: -1.15<br>Solvation: -0.91<br>Torsion: -0.89 | | | Alignment<br>Distortion |
| Human Coronavirus NL63 (Acc. No.: NC_005831.2:20472-24542) (Template: 6u7h.1A) | QMEAN: -0.63<br>Cβ: -0.10<br>All atoms: -0.68<br>Solvation: -0.93<br>Torsion: -0.28 | | | |
| Human coronavirus 229E (Acc. No.: NC_002645.1:20570-24091) (Template: 6u7h.1A) | QMEAN: -1.42<br>Cβ: -0.78<br>All atoms: -0.93<br>Solvation: -0.87<br>Torsion: -0.96 | | | |
| Human coronavirus OC43 strain ATCC VR-759 (Acc. No.: YP_009555241.1) (Template: 6nzk.1A) | QMEAN: -0.82<br>Cβ: -0.62<br>All atoms: -0.64<br>Solvation: -0.75<br>Torsion: -0.45 | | | Alignment<br>Distortion |
| Coronavirus (Acc. No.: NC_038294.1:21455-25516) (Template: MERS-CoV) | QMEAN: -0.54<br>Cβ: -1.21<br>All atoms: -1.84<br>Solvation: -1.45<br>Torsion: 0.19 | | | |
| Coronavirus (Acc. No.: NC_004718.3:21492-25259) (Template: SARS-CoV) | QMEAN: -2.82<br>Cβ: -0.65<br>All atoms: -1.83<br>Solvation: -1.84<br>Torsion: -2.01 | | | |
| SARS coronavirus 2 isolate Wuhan-Hu-1 (Acc. No.: NC_045512.2:21563-25384) (Template: COVID 19) | QMEAN: -3.63<br>Cβ: -0.98<br>All atoms: -2.08<br>Solvation: -1.99<br>Torsion: -2.69 | | | |

The system generated structures showed sequence identity with the homologous templates like Human coronavirus HKU1 and OC43 with template 6nzk.1A (Identity: 65.16% & 99.68% respectively), NL63 and 229E with 6u7h.1A (Identity: 65.22% & 99.10% respectively), MERS CoV with 5w9h.1.L (Identity: 99.69%), whereas, SARS CoV & COVID 19 with 6acc.1.A (Identity: 99.92% & 76.47% respectively). Structure quality assessment showed QMEAN values of two SARS strains were −2.82 and −3.63 for respective models (Table 1). This indicated the degree of native nature of the predicted structures in an universal scale [23]. Good quality structures were indicated with QMEAN Z-score closest to zero. QMEAN indicated the overall Z-score of Cβ, All atoms, Solvation and Torsion values. According to the Ramachandrans plot analysis on number of residues in favoured regions and in additionally allowed regions, human coronavirus HKU1 (98.5%), NL63 (99.2 %), 229E (99.8 %), OC43 (99.6 %), SARS CoV (99.8%), MERS CoV (98.3%) and COVID 19 (99.4%) were found as very good quality structures.

Higher degree of similarity between SARS CoV and SARS CoV-2 might indicate and help in the therapeutic and vaccination strategies with reference to the current global situation. However, an absolute higher degree of virulence and spreading nature of SARS CoV-2 is of great concern in the present scenario. Present prediction was further validated by the multiple sequence alignment of all CoVs and topology analysis of three spike glycoprotein structures of MERS CoV, SARS CoV and COVID-19 (Figure 1 and 2). Position specific multiple sequence alignment also showed the highest similarity of COVID 19 with the SARS CoV (Figure 1). Although having sequential diversity, all the selected spike glycoproteins showed some stretches of conserved sequences Figure 1. The position of N-terminal and C-terminal were found similar between SARS CoV and COVID-19 during topology analysis. On the other hand, a drastic difference was observed in MERS CoV in arrangement of secondary structures in the tertiary region (Figure 2).

**Fig 1.**
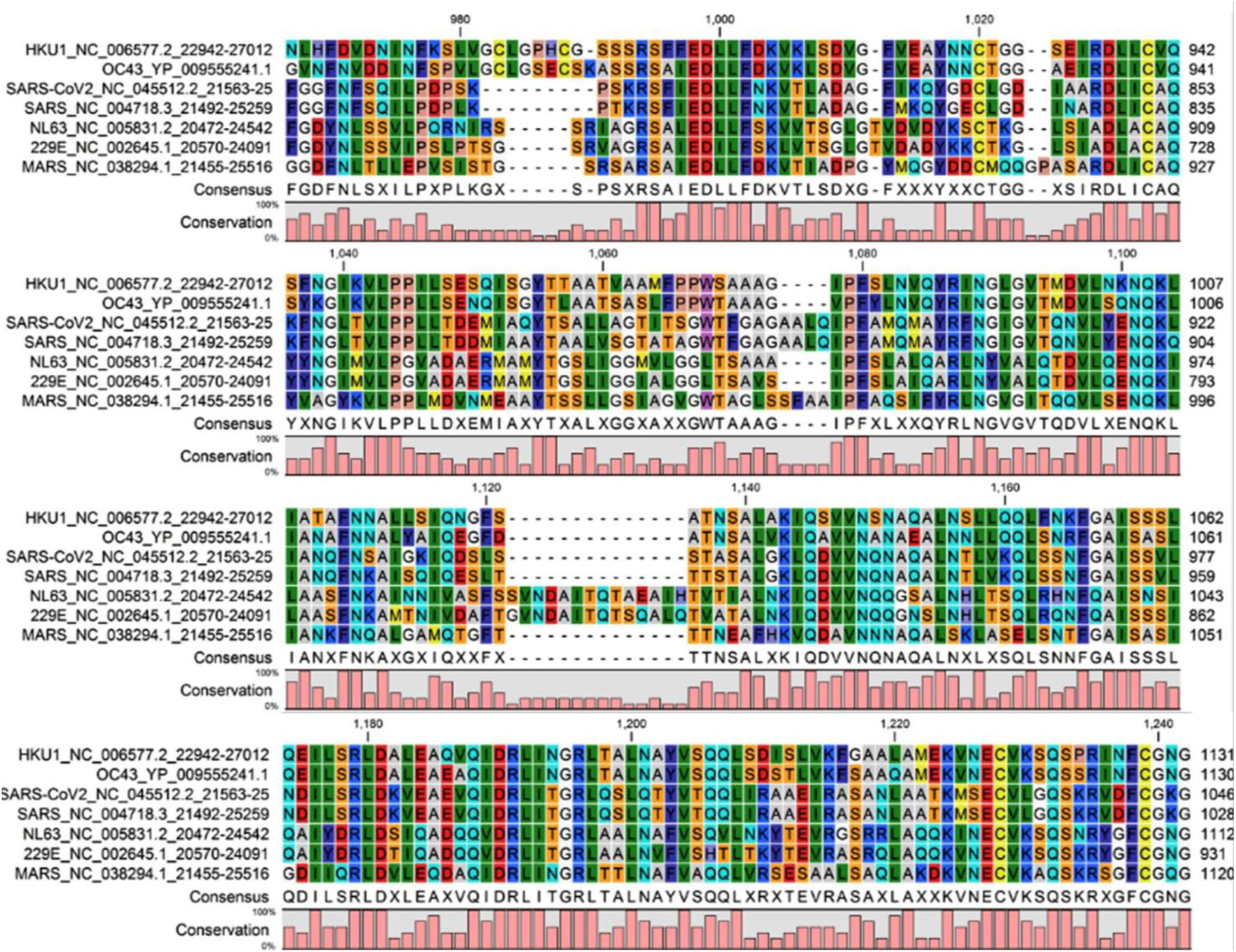
Multiple Sequence Alignment of selected coronavirus spike glycoproteins.

**Fig 2.**
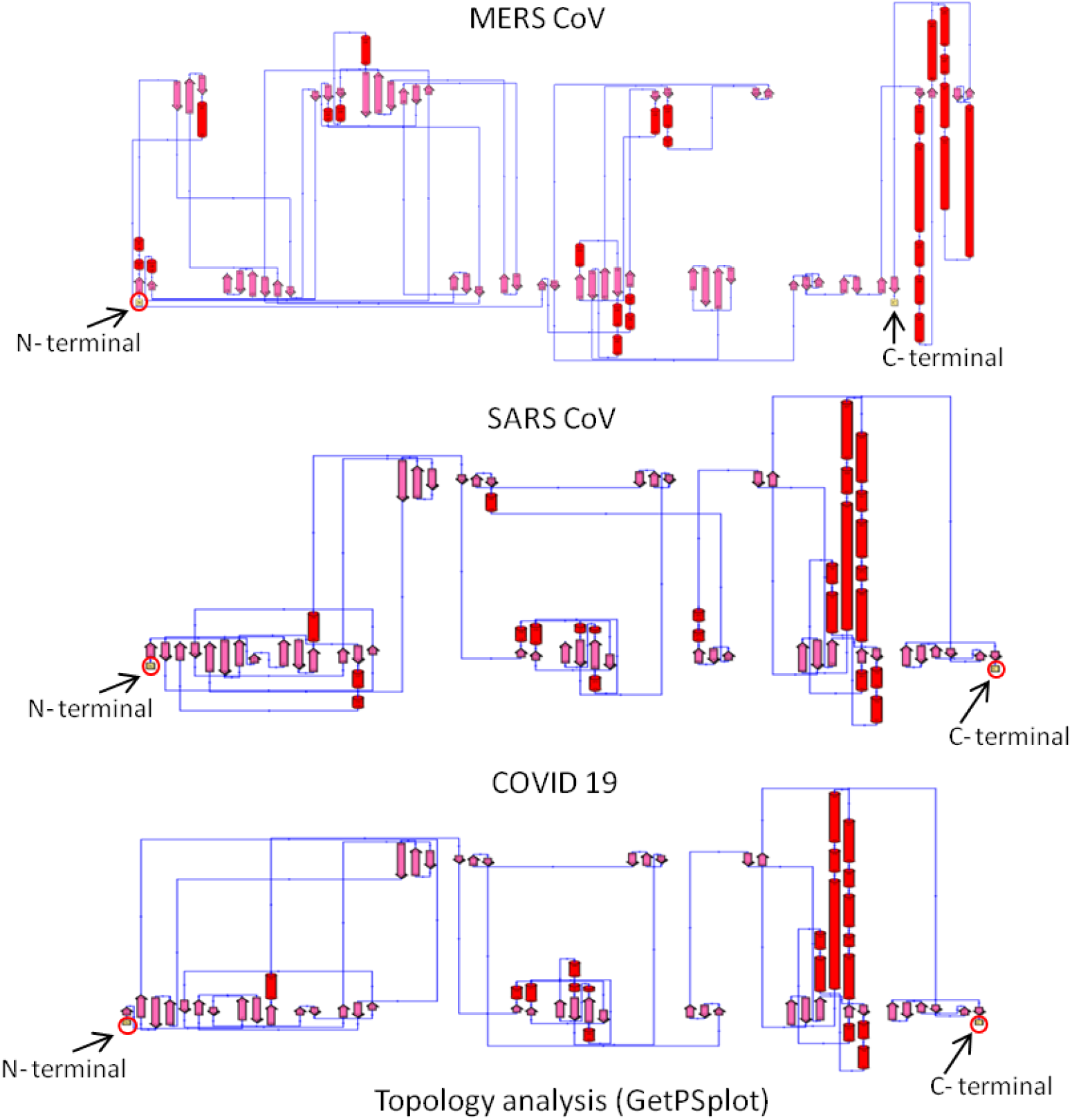
Topology analysis of three tertiary structures of MERS CoV, SARS CoV and COVID 19 spike glycoprotein.

### Conserved Motif Identification

Based on the alignment pattern, selected sequences were subjected to analysis different conserved motifs in the protein sequences. A total of 7 conserved motifs were analyzed (Figure 3). Most remarkably all the selected spike glycoprotein sequences were shown to have each 7 motif sequences in similar pattern and some of those conserved motif position were represented in figure 3. All the identified conserved motifs were individually subjected to protein BLAST for functional annotation. Where motif 1, 5, 6 and 7 showed similarity with spike protein of Human and Bat coronavirus origin, motif 2 shared similarities with spike structural protein of mouse coronavirus origin, motif 3 showed similarity with spike glycoprotein S from SARS CoV and motif 4 with spike glycoprotein from MERS CoV (Table 2). Highest percent identity of 91.84% was observed for motif 2. Functional gene ontology based conserved domain identification within 7 identified motifs were predicted against InterPro database. IPR002552 (CORONA_S2), PF01601 (CORONA_S2) domains were found within motif 1, 5 and 6. Similar domains were also observed in motif 2, 3 and 4 with an extra domain of SSF111474 (Coronavirus _S2 glycoprotein). No such domains were observed in motif 7. Identified domains were predicted with membrane fusion function (GO:0061025) and receptor-mediated virion attachment to host cell (GO:0046813). Whereas, those were detected as viral envelope (GO:0019031) and integral component of membrane (GO:0016021). Above analysis indicated that identified motifs were specific for coronavirus and they could be used as the markers for common coronavirus infection detection irrespective of COVID 19.

**Table 2.** Functional annotation of identified motifs and their functional gene ontology based conserved domain identification among all the selected corona virus spike proteins.

| Sl. No. | Motif Width | Function annotation through BLAST | Percent Identity in BLAST | Percent Identity with the Accession No. | Functional Gene Ontology based Domain Identification by Interproscan |
| --- | --- | --- | --- | --- | --- |
| 1 | 49 | spike protein [Human betacoronavirus 2c EMC/2012] | 75.51% | AGO06003.1 | IPR002552 (CORONA_S2), PF01601 (CORONA_S2) |
| 2 | 50 | spike structural protein [Longquan Aa mouse coronavirus] | 91.84% | AID16631.1 | IPR002552 (Corona_S2), PF01601 (CORONA_S2), SSF111474 (Coronavirus_S2 glycoprotein) |
| 3 | 41 | spike glycoprotein S [SARS coronavirus BJ182-4] | 85.00% | ACB69883.1 |  |
| 4 | 50 | spike glycoprotein [Middle East respiratory syndrome-related coronavirus] | 74.00% | AOR17480.1 |  |
| 5 | 50 | spike protein [Human coronavirus NL63] | 64.00% | AAY43188.1 | IPR002552 (Corona_S2), PF01601 (CORONA_S2), |
| 6 | 34 | spike protein [Bat SARS-like coronavirus] | 73.53% | AVP78031.1 |  |
| 7 | 25 | spike protein [Human coronavirus OC43] | 88.00% | AWW13559.1 | Not detected |

**Interproscan result:** Biological process involved: membrane fusion \(GO:0061025\), receptor-mediated virion attachment to host cell \(GO:0046813\)
Cellular components: viral envelope \(GO:0019031\), integral component of membrane \(GO:0016021\)

**Fig 3.**
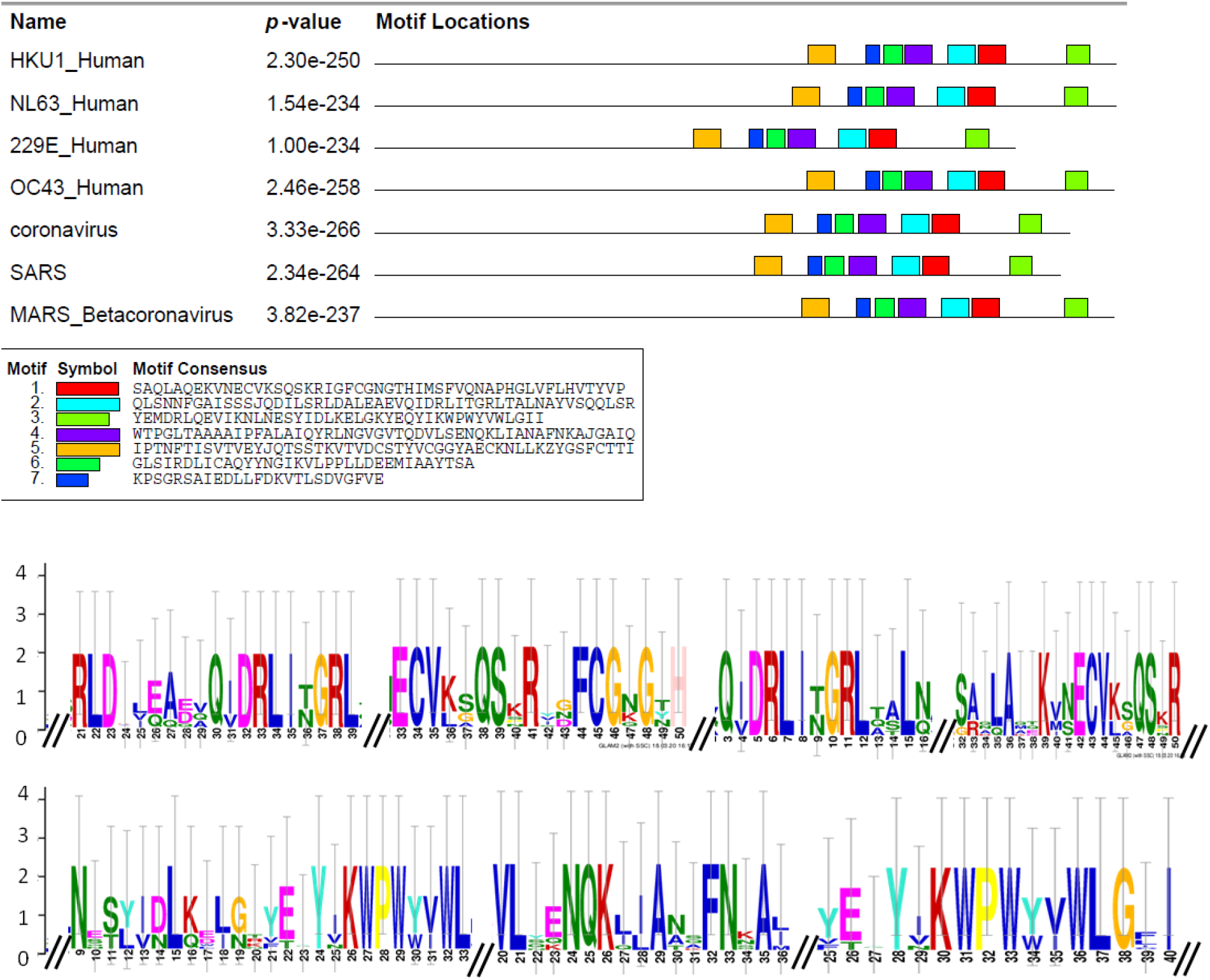
Conserved motif identification and their occurrence determination among all the selected corona virus spike proteins (upper panel). Representative portion of conserved Motif analysis data (lower panel).

### COVID 19 specific Epitope Designing

The epitope designing was conducted only with the COVID 19 spike glycoprotein sequence and structure. From the sequence analysis, 10 different locations were found which also showed similarity with SARS CoV and SARS COVID 2 spike glycoprotein in protein BLAST (Figure 4). Also the motif positions within the spike glycoprotein monomer were represented in figure 4. Epitopes 1, 4 and 5 were not represented in COVID 19 spike glycoproteins, as they were found to be embedded within the virus envelop. Among the others, motifs 2, 3, 6 and 7 were found at the interior location of spike glycoprotein monomer but motifs 8, 9 and 10 were found at the surface of the structure, which could be used as the immunological targets for the proper diagnosis and treatment of COVID 19. The important issue of epitope finalization could be confronted by the factor of possible transition between pre-fusion and postfusion spike structural distortion. Specific mutant structure has been designed and tested to be resistant to conformational change after ACE2 binding and protease cleavage at the S1/S2 site [34]. This may be indicative to searching suitable epitope which may remain unhindered from pre-to post-fusion state transition.

**Fig 4.**
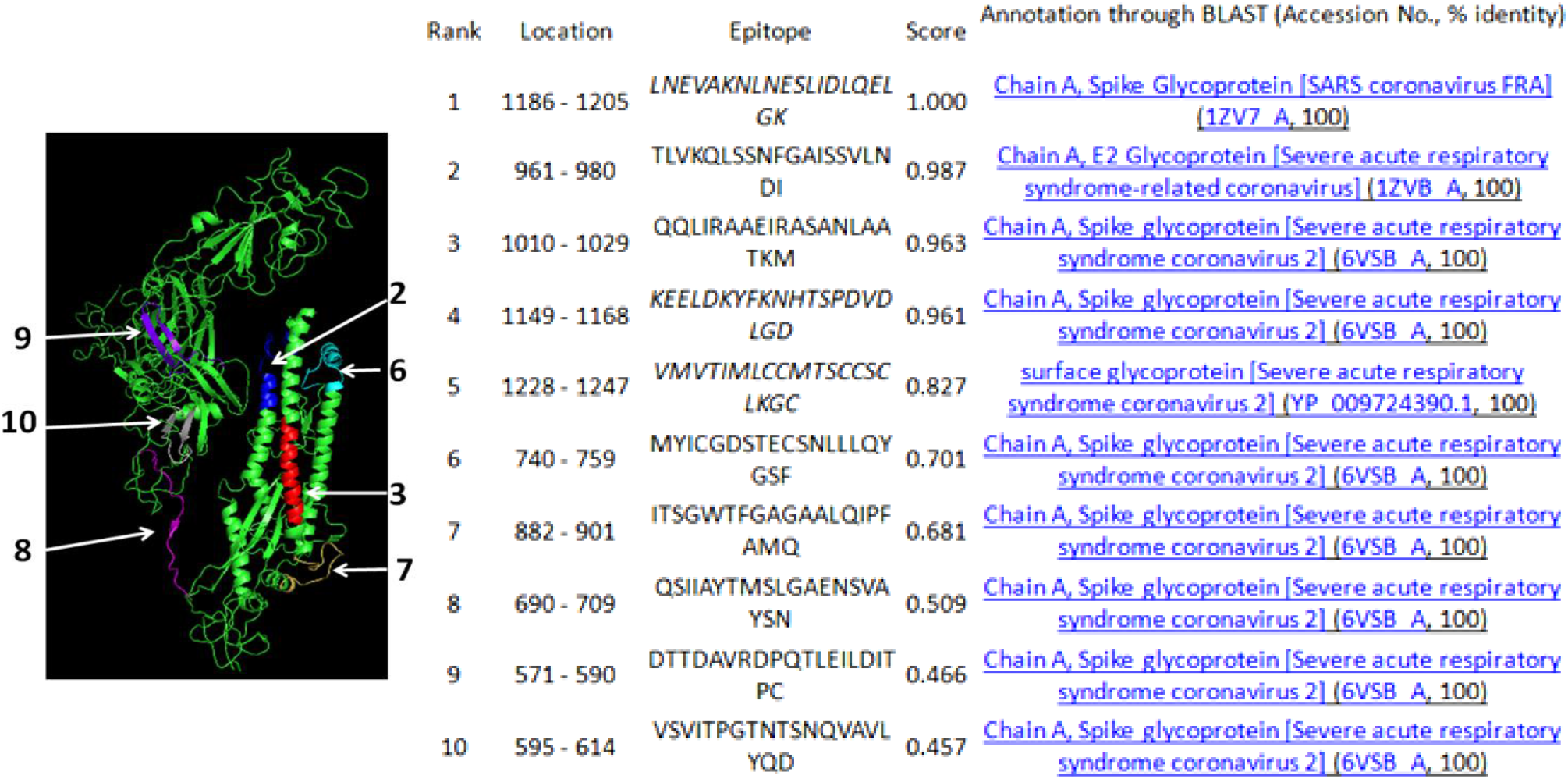
Epitope predicted inside COVID 19 spike glycoproteins

### Analysis of epitope binding to specific MHC Class II

The proper type of Major Histocompatibility Complex (MHC) selection for identified COVID 19 epitope was performed and enlisted in Table 3. All the epitopes were individually screened against 5 DPA, 6 DQA and 662 DRB alleles from MHC class II for best fit analysis. As, we have analyzed the spike glycoprotein of COVID 19 which is an infectious particle, transmit from one infected individual to another, alleles of MHC class II were selected for viral epitope specificity analysis. Where HLA-DRB1*01:13 was observed to bind with 2B, 3B, 4B, 5B, 7B and 8B epitope sequences identified on the basis of IC50 value. Among them sequence IIAYTMSLGAENSVA (epitope 8B) was shown with lowest IC50 value of 7.11. On the other hand, HLA-DRB1*04:04 was found for both the sequence of TIMLCCMTSCCSCLK (epitope 5A) and SIIAYTMSLGAENSV (epitope 8A) on highest consensus percentile rank basis. Highest value of 9.5 was found for motif 5. Similarly HLA-DRB1*04:08 was observed for the sequences VRDPQTLEILDITPC (epitope 9A) with highest 9.50 and VSVITPGTNTSNQVA (epitope 10A) with 7.90 consensus percentile rank value. Individual MHC class II molecules were found for others (Table 3). The threshold value of highest consensus percentile rank was selected as 10 for all. As a whole, highest Consensus percentile rank value of 10 was observed for sequence QQLIRAAEIRASANL (epitope 3A) and lowest IC50 value of 7.11 was observed for sequence IIAYTMSLGAENSVA (epitope 8B).

**Table 3:**
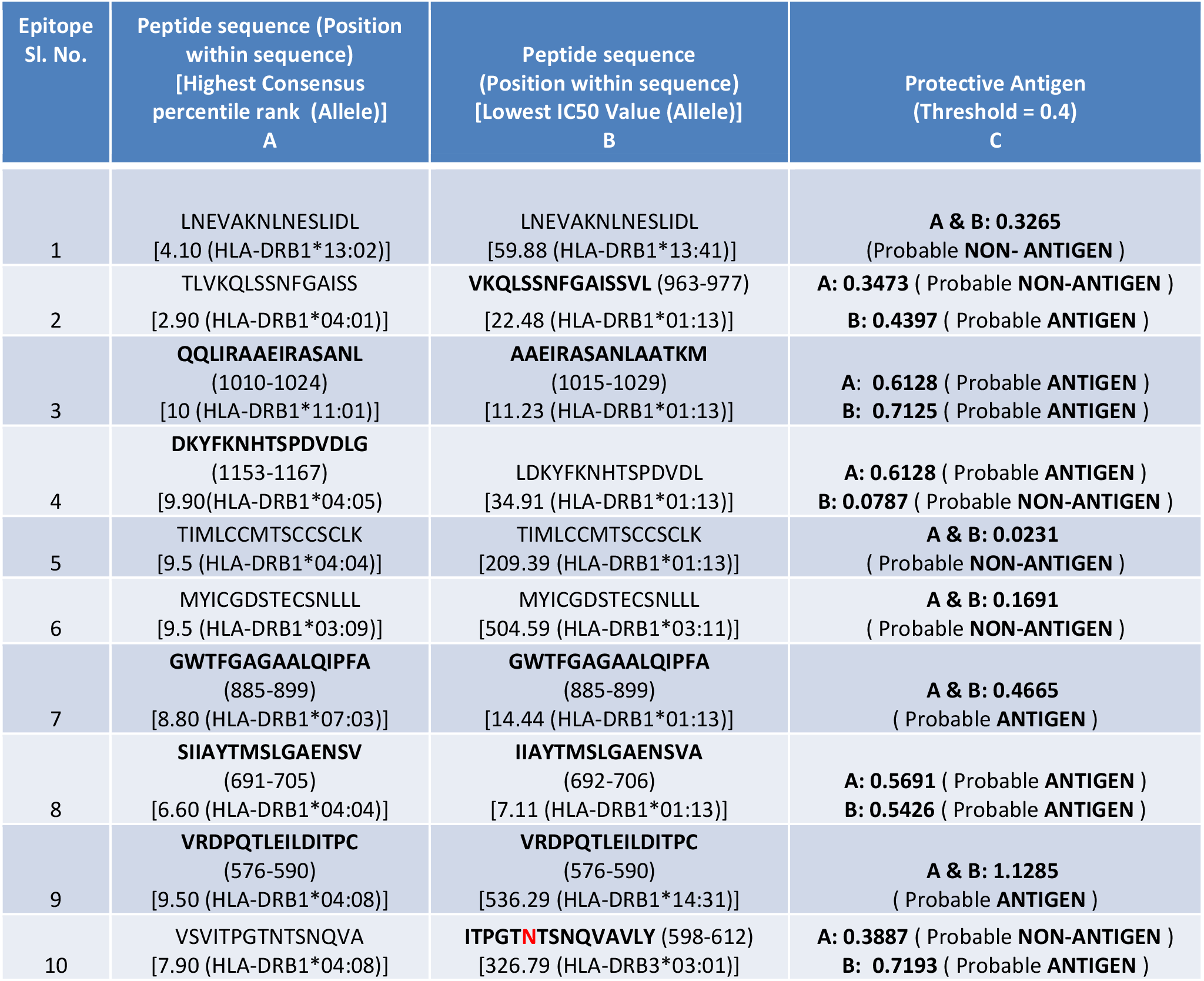
Coronavirus 19 spike protein epitop analysis for best MHC class I allele selection on the basis of Highest Consensus percentile rank and Lowest IC50 Value (A & B). Determination of antigenic property of identified epitops (C).

The antigenic property of identified target sequences from epitopes was also predicted on the basis of threshold value of 0.4. Below the threshold value, the sequence has been considered as non-antigenic and sequences with above value were antigenic in nature. A total of 9 antigenic sequences were detected (Table 3), among them two sequences AAEIRASANLAATKM (epitope 3B) and ITPGTNTSNQVAVLY (epitope 10B) were found with higher threshold value of 0.7125 and 0.7193 respectively. As the location of 3B was more interior 10B could be used as potent antigen. According to epitope locations (Figure 4) and antigenic nature, other sequences like 8A&B, 9A&B could be the target also.

### Glycosylation and structural modification

Coronavirus spike proteins are glycosylated in nature where N-acetyl glucosamine (NAG) is the main component. Glycan shielding and possible epitope masking of an HCoV-NL63 has been observed which may be the barrier for proper immunogenic responses [8]. Comparative analysis between glycosylated and non-glycosylated protein revealed some structural modification at the epitope locations. Among the identified epitopes 10B with sequence ITPGTNTSNQVAVLY (598-612) was found with N linked glycosylation at 603 position. The structural modification of this epitope was analyzed using non-glycosylated protein structure of COVID 19 (Acc. No.: NC_045512.2:21563-25384) and gltcosylated COVID 19 protein (PDB ID: 6vsb). Effect of glycosylation on protein structures revealed that glycosylated conformation was more organised (Figure 5a) than non-glycosylated one (Figure 5b). Secondary structural comparison between two epitopes showed more organised structure with attached NAG residue (Figure 5d) whereas a shorter β-sheet structure was observed when NAG is removed from the structure (Figure 5c). The peptide interactive site of 10B epitope was blocked due to NAG attachment. As a result of which antibody binding to the antigen may hamper. The NAG residue directly binds with N or ASN amino acid residue (Figure 5e). So the removal of NAG from the spike glycoprotein structure is difficult. Structural distortion between glycosylated and non-glycosylated epitope 10B at tertiary level indicated that removal of NAG may distort the structure of epitope (Figure 5f). Again that may hamper the proper antigenantibody binding.

**Fig 5.**
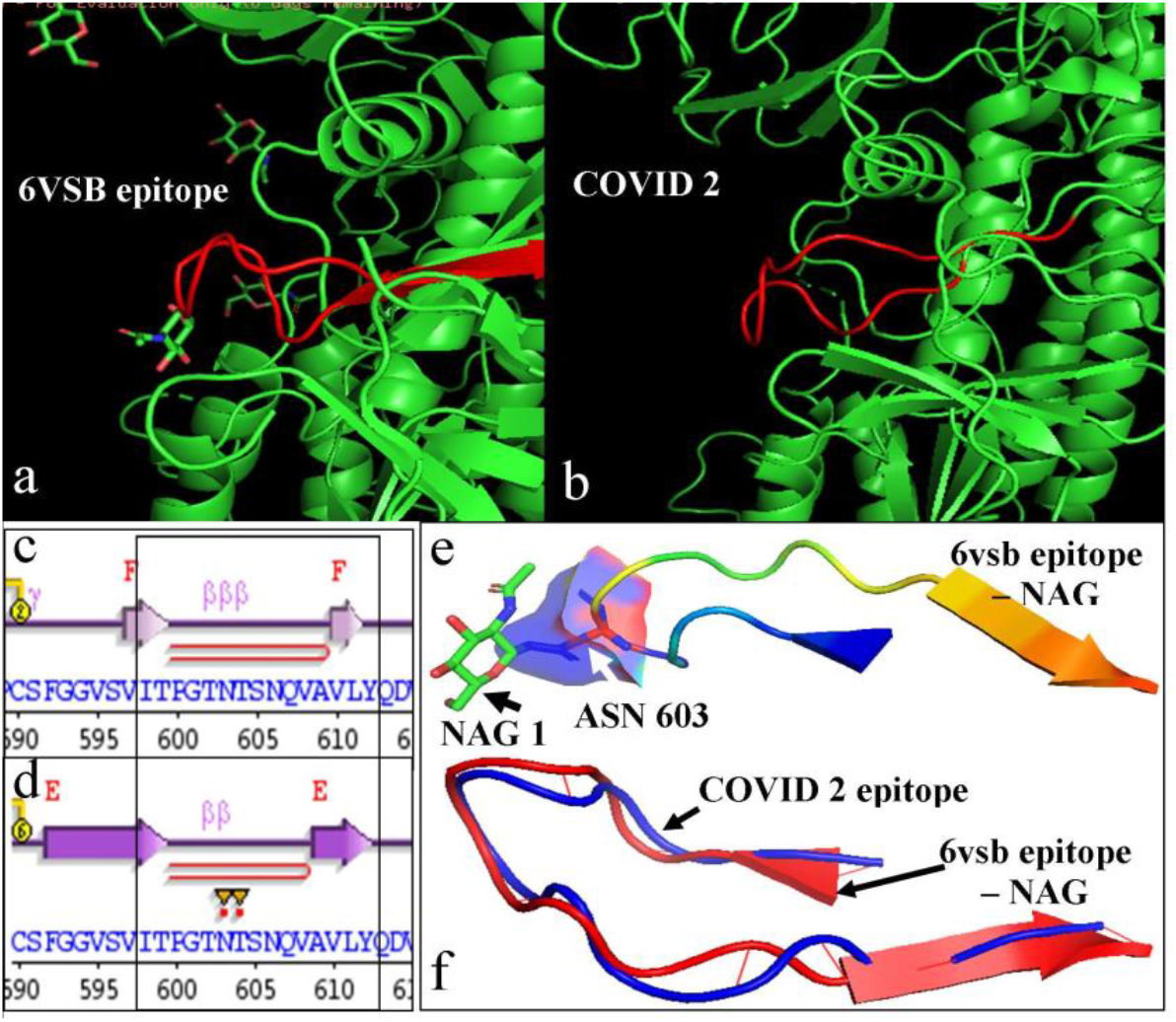
Effect of glycosylation on protein structure. 10B epitope position on COVID 19 spike protein (PDB ID: 6vsb), NAG attached with N residue at the 603 position (a). 10B epitope position on COVID 2 or COVID 19 spike protein (Acc. No.: NC_045512.2:21563-25384), no NAG attached with N residue at the 603 position (b). Secondary structure of epitope 10B without NAG attachment (c) and with NAG attachment (d). Close view of NAG attachment with N residue in 6vsb at position 603 (e). Structural alignment between glycosylated and non-glycosylated 10B epitope structure (f).

### Effect of epitope glycosylation on MHC class II – epitope binding

In this section energetics of epitope attachment with MHC class II HLA-DRA, DRB was determined in presence and absence of NAG at the 10B epitope structure through molecular docking (Figure 6). Docking results showed that without NAG, the binding efficiency of 10B epitope at the epitope binding site of MHC class II HLA-DRA, DRB molecule was very high. Among the 10 best docking posture, 1, 2, 3, 7 & 10 were found at the desired location with docking score of −3552.23, −3472.43, −3436.90, −2767.44 and – 2185.81. Whereas, tertiary structure of epitope 10B with NAG revealed less affinity to MHC class II HLA-DRA, DRB molecule. Only 3 postures, 5, 7 & 10 were found at the desired position with docking score of - 3085.38, −2949.73 and −2141.10. The best docking of posture 1 (without NAG) and posture 5 (with NAG) were represented in Figure 6 where amino acid attachment differences were clearly indicated in Figure 6b and 6e. Like the docking score of posture 1 (without NAG) −3552.23, it also showed the binding free energy of complex, −36.97 (kcal/mol). Whereas, docking score of posture 5 (with NAG) −3085.38, showed binding free energy of complex, −30.06. That indicated the rigid binding of 10B epitope when it lacks the NAG molecule. The interactive analysis also revealed that without NAG, 10B binds with more amino acids of MHC class II HLA-DRA, DRB (Figure 6c) where structural stabilization by hydrogen bond networking was noticed. But, the bindings were less when NAG residue was attached (Figure 6f). This result indicated that the attachment of NAG with epitope also made it difficult for MHC class II molecules to proper representation of epitope.

**Fig 6.**
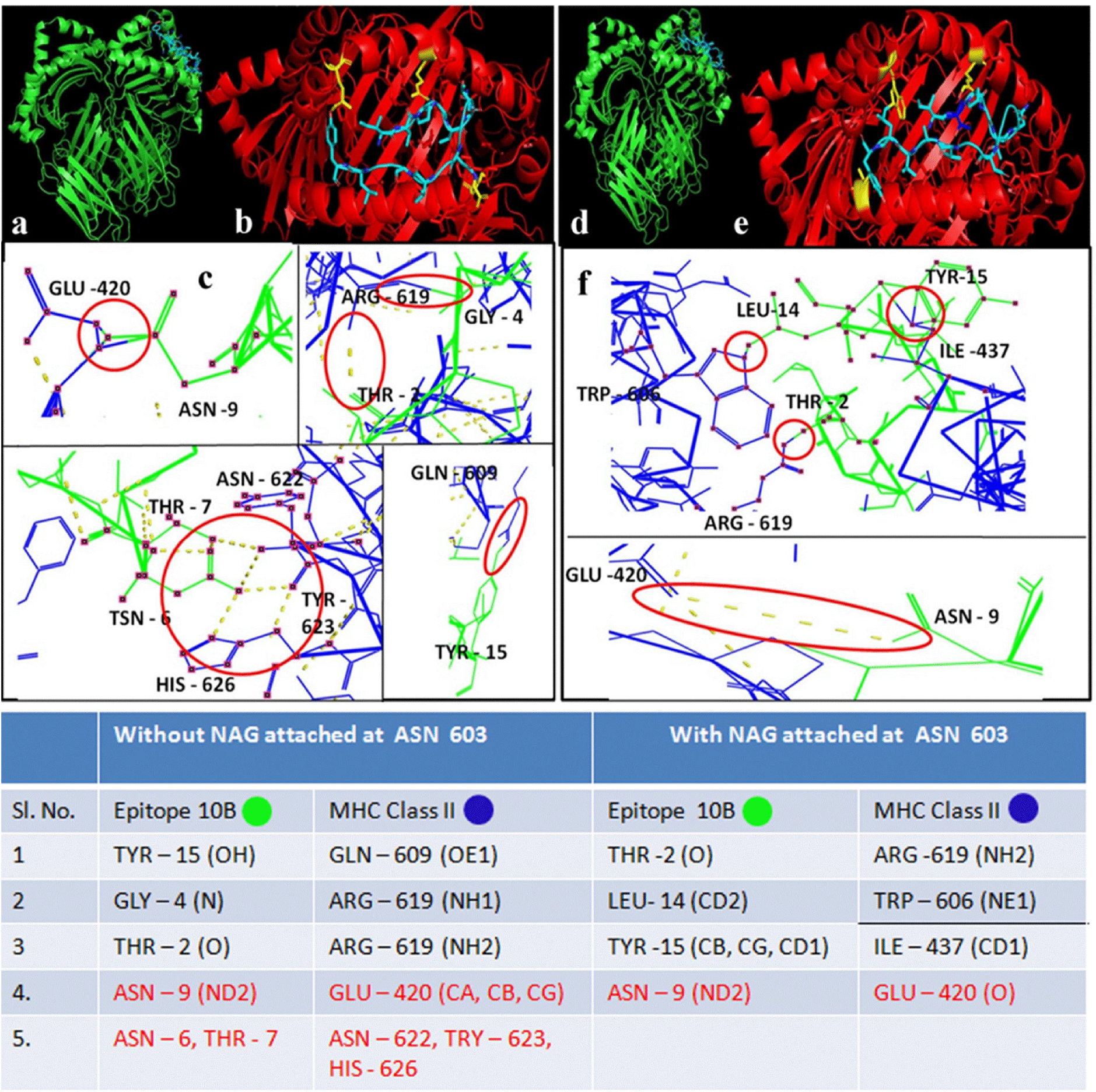
Effect of epitope glycosylation on MHC class II – epitope 10B binding. Without NAG epitope 10B binding to MHC class II HLA-DRA, DRB epitope binding site (a, b) and different molecular interactions of 10B epitope with MHC class II (c). With NAG epitope 10B binding to MHC class II HLA-DRA, DRB epitope binding site (c, d) and different molecular interactions of 10B epitope with MHC class II (f). Lower panel of tabulated image describes amino acids responsible for stable binding between epitope 10B and MHC molecule in presence and absence of NAG.

Though, epitope 8 A & B were also present at the surface of the spike glycoprotein but was found to wrapped with a short segment IGAEHVNNSYECD (651-663) carrying a glycosylation at N residue position 657 (Figure 7a). As a result of which antibody accessibility to this epitope may also be difficult. Whereas, surface epitope 9 with sequence VRDPQTLEILDITPC (576-590) showed highest antigenecity of 1.1285 (Table 3) and highest consensus percentile rank of 9.50 and found free of any direct or indirect NAG attachment pattern (Figure 7b). On that basis, it was further analyzed for MHC II HLA-DRB1 (PDB ID: 5jlz) binding through molecular docking. Among the best 10 docking posture, 8 were found at the desired position of MHC molecule (Figure 7c). The best position 1 was represented in figure 7 d&e. A rigid interaction with six amino acids and one hydrogen bonded amino acid of MHC molecule was detected for proper representation of epitope 9. So, this could be a target for COVID-19 vaccine development.

**Fig 7.**
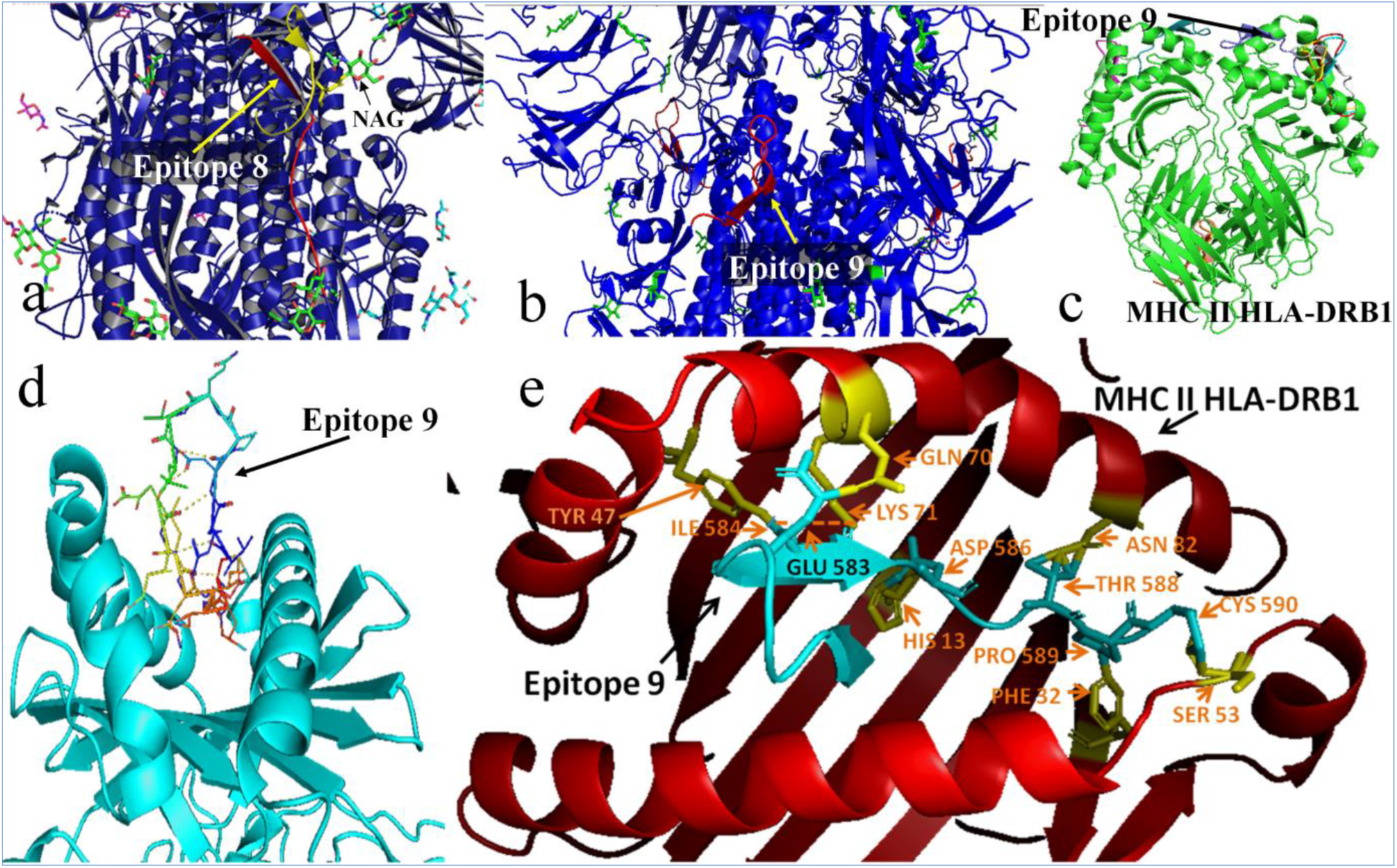
Position of Epitope 8 in COVID 19 spike protein with indirect NAG masking, PDB ID: 6svb (a). Position of Epitope 9 in COVID 19 spike protein with no direct or indirect NAG attachment, PDB ID: 6svb (b). Among 10 best docking positions, 8 were found in epitope presenting site of MHC II HLA-DRB1, PDB ID: 5jlz (c). The best docking posture of epitope 9 with MHC II HLA-DRB1 (d) and its specific interaction pattern (e).

Modifications in spike proteins structure during receptor mediated host cell entry and further prediction on post-fusion events may result in success in vaccination strategies or blocking entry. Host protease processing during viral entry and how different lineage B viruses can recombine to gain entry into human cells are the points also to be noted [35]. SARS CoV-2 induced severe and often lethal lung failure is caused due to its inhibition of ACE-2 expression [36]. So, keeping the ACE-2 normal functioning but blocking viral entry is the most challenging issue right now. Possible suitable epitope as screened in the current study may be helpful in this global pandemic situation. The history of last two decades’ outbreak of these types of virus is very much evident. The present situation justifies further advanced studies with proper infrastructure and fund-resources facilities at a global scale to eradicate current or any possible future outbreak.

## Declarations

- Ethics approval and consent to participate: N/A
- Consent for publication: Yes
- Availability of data and material: Yes
- Competing interests: None
- Funding: Institutional
- Authors’ contributions: Concept-SM and AM, Study design-SM and AM, Experiments-AM and DS, Analysis-all three authors, Manuscript writing-SM and AM, Revision-all authors
- Acknowledgements: OIST members

